# Examining urban weed response to elevated salt levels

**DOI:** 10.64898/2026.11.05.763933

**Authors:** Maximilian M. Carroll, Jennifer M. Cocciardi, Ava Hoffman, Kelsey Coates, Eric G. Yee, Meghan L. Avolio

**Author notes:** Corresponding Author: Maximilian M. Carroll.

## Abstract

Stressors from the urban environment cause adverse effects on city-dwelling organisms. While there is evidence that some species are adapting to urban stressors, such as the urban heat island and impervious surfaces, other stressors such as road salt are less studied. We investigated whether plants are adapting to elevated salt concentrations with two common garden experiments using three common weed species: *Veronica arvensis*, *Stellaria media*, and *Lamium purpureum*. We collected plants from parks around Baltimore, MD, USA, collecting plant maternal lineages from roadside soils (i.e. higher soil salinity) and inner park soils (i.e. lower soil salinity). We hypothesized that roadside lineages would tolerate salt treatment better than park lineages. In our germination experiment on wild-collected seeds, we observed a non-significant trend where the germination rate of roadside *L. purpureum* lineages was reduced less by salt treatment than for park genetic lineages, while *S. media* had similar declines between groups. In our greenhouse experiment on the F1 generation, roadside *V. arvensis* lineages displayed a less adverse reaction to salt treatment than park *V. arvensis* lineages. *L. purpureum* did not display this difference, but roadside lineages of both species exhibited less tissue damage overall. The results of our experiments were likely influenced by the maternal environment, and work on F2 generations is required to confirm them. Our findings show some support for urban salt tolerance adaptation, and point to the heterogeneity of the urban environment and the complexity of plant reactions to salinity.

## Introduction

Plants in cities face unique environmental stressors, including habitat fragmentation (Cheptou, et al., 2008), increased temperatures (Woudstra et al., 2024), and elevated salt levels (Equiza et al., 2017). There is growing evidence that plants are undergoing adaptive evolution in response to these stresses (Lambert et al., 2021; Thompson et al., 2016; Yakub & Tiffin, 2019; Yee, 2023). As cities continue to expand, further understanding plant response to urban stressors is essential for preserving the ecological functions that support urban life. A multitude of traits have been observed to diverge between urban and rural plant populations, including seed dispersal in holy hawksbeard (*Crepis sancta*) (Cheptou, et al., 2008), flowering time in common ragweed (*Ambrosia artemisiifolia*) (Gorton et al., 2018), and cyanogenesis in white clover (*Trifolium repens*) (Thompson et al., 2016). Although adaptive trends are emerging across cities, the strengths of such trends can vary considerably (Santangelo et al., 2022a), and the reactions of plant populations to urban stressors can vary across cities and species (Yee, 2023). Since these patterns of adaptation can vary by species (Hoffman et al., 2026), there is a need for comparative studies across species to determine if patterns of urban adaptation in plants are generalizable (Santangelo et al., 2020).

Winter salt application in cities is widespread, with up to 24 million tons of salt applied annually to roads across the United States (Tiwari & Rachlin, 2018). The application of road salt increases sodium concentrations in roadside soils and affects water quality in urban streams (Daley et al., 2017). High salt concentrations can impair plant growth, disrupt physiological processes, and, in extreme cases, prove lethal (Łuczak et al., 2021). Due to the severity and variability of plant reactions to salt stress (Munns & Tester, 2008), salt tolerance has been identified as a consideration for selecting turfgrasses (Friell et al., 2012) and urban trees (Šerá, 2017). Though salt is identified as a major stressor for plants in cities, most literature on salt tolerance focuses on an agricultural context (e.g. Chartzoulakis & Klapaki, 2000; Houshmand et al., 2005). Plant species vary considerably in their ability to tolerate salt (Munns & Tester, 2008). A study of salt tolerance in weedy species found that roadside populations of one weed (*Dipsacus fullonum* subsp. *sylvestris*) exhibited elevated salt tolerance compared to non-roadside populations while two other weeds (*Asclepias syriaca* and *Hesperis matronalis*) did not (Beaton & Dudley, 2004). Further investigation revealed that both maternal effects and genetic influence contributed to observed differences in salt tolerance (Beaton & Dudley, 2007). Salt tolerance in roadside populations may thus be influenced by both environmental and genetic factors, including local adaptation, developmental plasticity, and maternal effects. These studies demonstrate the complexity of plant reactions to elevated salinity, creating a need for research across multiple species to better understand the responses of roadside plants to salt stress.

In addition to unique stressors, the urban environment is characterized by high levels of heterogeneity compared to other environments. Broad geographic categorizations, such as urban or rural, can miss considerable variation due to land use, impervious surface cover, and other factors changing on a small spatial scale (Cadenasso et al., 2007). For example, impervious surface cover, population density, vegetation cover, and topography all contribute to small-scale urban heat island variation (Mokhtari et al., 2022). While the urban-rural gradient is a common and powerful paradigm in urban ecology to study adaptation (McDonnell & Pickett, 1990), it cannot account for spatial heterogeneity within cities, which is particularly important for sessile organisms such as plants. Many existing studies take characteristics of individual sites into account alongside the urban-rural gradient (e.g. Song et al., 2019), but directly comparing nearby subhabitats within an urban environment is less common. Road salt application, in particular, varies greatly over short distances, with its negative effects concentrated on roadsides (Coates et al., 2025; Łuczak et al., 2021). The high variability of road salt concentration across adjacent soils provides an opportunity to study differences in plant responses to salt stress across small spatial scales.

This study aimed to evaluate whether three common urban weeds—chickweed (*Stellaria media*, Caryophyllaceae), red dead-nettle (*Lamium purpureum*, Lamiaceae), and corn speedwell (*Veronica arvensis*, Plantaginaceae)—are adapting to tolerate roadside elevated salt content. All three species are commonly found throughout urban areas (Table 1), and are exposed to high soil salinity along roadsides, making them suitable for studying salt tolerance adaptations. To address our overarching question, we conducted two experiments, a seed germination experiment and a greenhouse experiment, on the F1 generation. For both experiments, we collected seeds of focal species from co-located roadside and park soils, and tested whether park-collected lineages and roadside-collected lineages differed in salt tolerance. We hypothesized that across experiments and species, roadside-collected lineages would better tolerate salt compared to park-collected lineages, due to higher salt exposure and therefore greater selection for salt tolerance, or due to maternal effects priming offspring to tolerate salt.

**Table 1:** Information on our focal species. All three species are annual non-native plants. Photographs are from Wikimedia Commons. *S. media* photo credit: AnRo0002, own work, CC0, https://commons.wikimedia.org/w/index.php?curid=102504142. *L. purpureum* photo credit: AnRo0002, own work, CC0, https://commons.wikimedia.org/w/index.php?curid=32001155. *V. arvensis* photo credit: AnRo0002, own work, CC0, https://commons.wikimedia.org/w/index.php?curid=58337797.

| Species (common and scientific names, and family) | Lifespan | Reproduction | Phenology | Photograph |
|---|---|---|---|---|
| Chickweed (*Stellaria media,* Caryophyllaceae) | Winter annual in temperate areas (DeFelice, 2004) | High self-pollination rates, some insect-mediated cross-pollination (DeFelice, 2004) | Germinates in late fall or early spring; flowers year-round under moderate conditions (Turkington et al., 1980) |  |
| Red dead-nettle (*Lamium purpureum*, Laminaceae) | Winter annual in temperate areas (Baskin & Baskin, 1984) | Self-pollinates, facultatively insect-pollinated (Macior, 1978) | Germinates in autumn; flowers and sets seed by early summer (Baskin & Baskin, 1984) |  |
| Corn speedwell (*Veronica arvensis*, Plantaginaceae) | Winter annual in temperate areas (Baskin & Baskin, 1983) | Often self-pollinates, facultatively insect-pollinated (Boutin & Harper, 1991) | Germinates in mid-late autumn; flowers in late spring or early summer (Baskin & Baskin, 1983) |  |

## Methods

### Study location

Seeds for this study were collected from parks in and around the urban area of Baltimore, MD, USA. Baltimore has a humid subtropical climate (Köppen: Cfa), within the broader temperate climate classification (Beck et al., 2018), and is located in USDA Plant Hardiness Zones 7b and 8a (USDA, 2023). Baltimore’s summers are hot and humid, with a mean July temperature of 25.9 °C (78.6 °F) and mean July precipitation of 12.65 cm (4.98 in) (NWS, 2026). Snow accompanies Baltimore’s winters, with average January temperatures of 1.2 °C (34.2 °F) and average total January snowfall of 17.27 cm (6.8 in) (NWS, 2026). Data on the application of road salt in Baltimore City are not publicly available, but the surrounding Baltimore County applies an average of 60,789 tons of road salt annually (Androutsopoulos et al., 2024). Application of road salt in Baltimore City and the surrounding Baltimore County drives plumes of NaCl-contaminated groundwater tied to winter snowfall, indicating that road salt application runs off roads and through soils (Snodgrass et al., 2017) with a strong seasonal signal during winter in the Mid-Atlantic (Moore et al., 2020).

### Species description

*S. media*, *L. purpureum*, and *V. arvensis* (Table 1) are annual flowering plants native to Eurasia which have spread widely in North America (Baskin & Baskin, 1983, 1984; DeFelice, 2004). These species grow and reproduce quickly, are relatively short, and are classified as weeds. In subtropical climates such as Baltimore, the three species behave primarily as winter annuals, where they germinate in fall, overwinter as seedlings or rosettes, and produce flowers in spring. *S. media* produces small white flowers throughout the year under favorable conditions, generally spring and summer in temperate climates, and sets seed for two to three months after flowering (DeFelice, 2004). *L. purpureum* produces pink axillary flowers in early spring and sets seed by early summer (Baskin et al., 1986; Baskin & Baskin, 1984). *V. arvensis* produces blue-lavender flowers starting in mid-spring, and sets seed in late spring or early summer (Baskin & Baskin, 1983). The three species are each capable of self-pollination and insect pollination (Boutin & Harper, 1991; DeFelice, 2004; Macior, 1978). The autumn germination of the three species means they are likely to be exposed to road salt within their life cycle, before spring rains wash it out of the soil. All three species are common along Baltimore’s roadsides, resulting in exposure to elevated salinity (Coates et al., 2025), and are often unmanaged, allowing them to grow and reproduce with little direct human selection.

### Sampling design

We sampled at eight parks across Baltimore (Fig. 1; Tables S1 and S2), hereafter a collection site. At each collection site, we sampled from the roadside collection environment and from the park collection environment. We defined the roadside collection environment as lawn within one meter of the street, and the park collection environment as lawn at least 20 m away from any street or impervious surface. Thus, for each collection site, we had two collection environments (roadside or park). Within each collection environment, all maternal lines of a species were collected at least 10 m away from each other.

**Figure 1:**
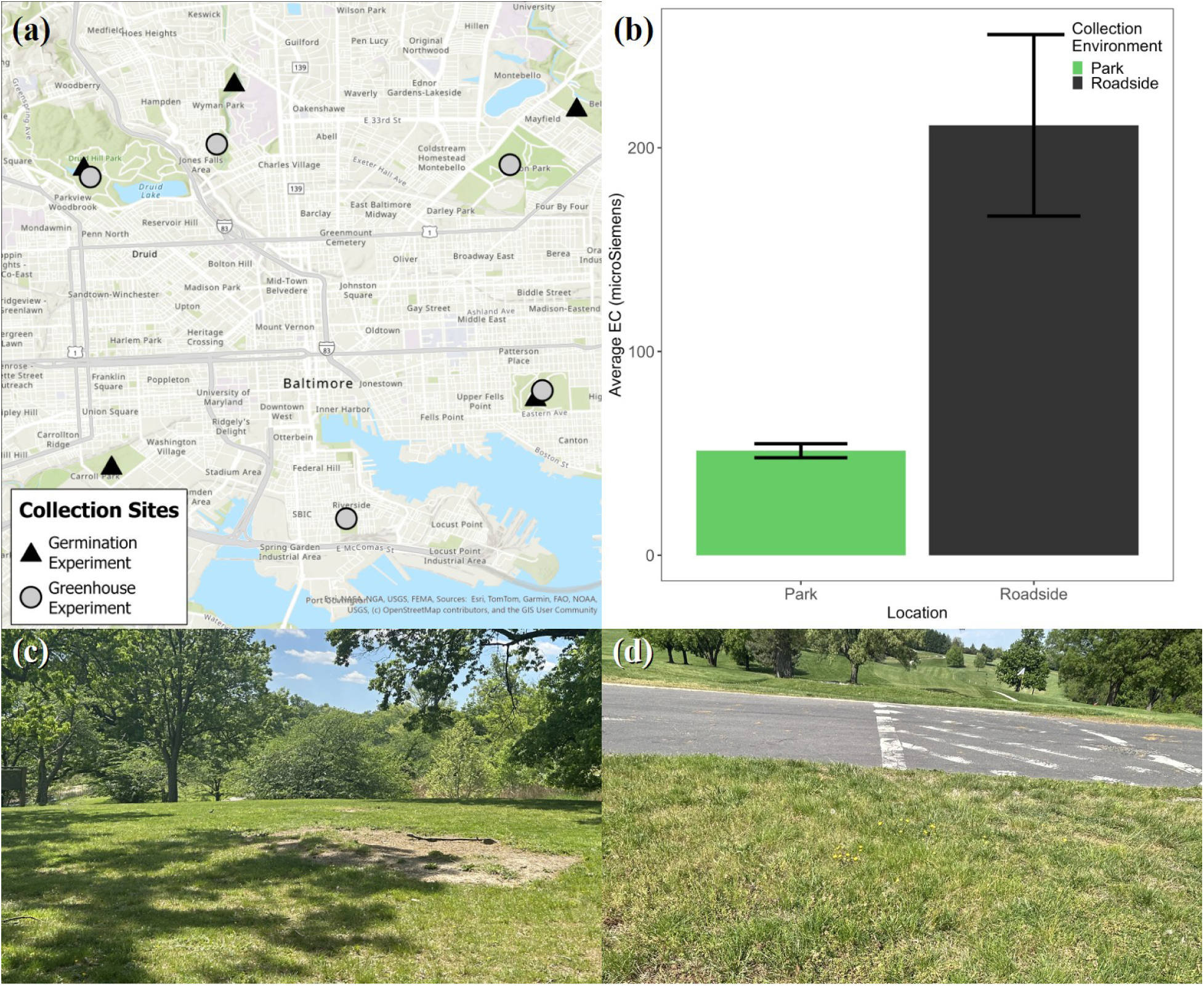
(a) Seed collection sites for our study. Black triangles represent where seeds were collected for our germination experiment, and light gray circles represent where seeds were collected for the greenhouse experiment. At each park, we collected plants from both park and roadside collection environments. (b) Average electrical conductivity (EC) of park collection environments and roadside collection environments. Error bars represent standard error. Photos depict (c) a representative park collection environment and (d) a representative roadside collection environment. Roadside and park photo credit: Maximilian M. Carroll.

### Soil sampling

We collected soil samples alongside our plant samples to confirm differences in soil salinity between our park and roadside collection environments (Table S3). Soil samples of 10 cm depth were collected at six points per site, with three samples from the roadside collection environment and three samples from the park collection environment. Initial sampling occurred in April and May 2024 alongside seed collection, but due to equipment error, resampling was required. Soil resampling occurred in April 2025 in the same manner as described above, but with six samples from the roadside and six samples from the inner park, resulting in 58 samples collected in total (two samples were discarded due to excess rock content; Table S3). Though road salt application varies annually with winter snowfall (Snodgrass et al., 2017), total snowfall in Baltimore was 28.7 cm (11.3 in) for the winter of 2023-2024, compared to 32.3 cm (12.7 in) for the 2024-2025 winter (NWS, 2026), so we presume our 2025 soil samples are representative of saline roadside conditions for our experimental plants. Roots and rocks were separated from the soil, which was then ground into a fine powder using a mortar and pestle. To measure electrical conductivity, a proxy for soil salinity (Rhoades et al., 1990), 10 g of soil were mixed with 50 mL of distilled water, stirred for two minutes using a milk frother, and left to settle for five minutes. Electrical conductivity was then measured using an Oakton EcoTestr™ EC1 Waterproof Conductivity Pocket Tester (Cole-Parmer Instrument Company, Vernon Hills, Illinois).

### Germination experiment

Seeds for *S. media* and *L. purpureum* were collected from parent plants, hereafter maternal lines, from five collection sites in Baltimore (Figure 1; Table S1). We collected up to 10 maternal lines of each species from each collection environment within each collection site in April and May of 2020. Seeds were separated from their parent plants, then cold stratified for 20 months at 4 °C before germination. Seeds were placed in plastic petri dishes with a diameter of 7.62 centimeters (3 inches) on a paper towel medium, which was moistened with deionized water as needed. We only included maternal lines that produced at least 20 seeds, to allow for five seeds per treatment per line (Table S1). Each dish was covered with Saran wrap, and placed in a CARON model 7305-22-1 growth chamber (Caron Scientific, Marietta, Ohio). *S. media* and *L. purpureum* were grown with a 12 hour photoperiod at 17 °C at 60% relative humidity (Baskin & Baskin, 1984; Roberts & Lockett, 1975).

We conducted a 2 × 4 fully factorial experimental design with two collection environments (roadside or park) and four salt treatments: control (no salt), low (75mM NaCl), medium (200 mM NaCl), and high (400 mM NaCl) based on a salt tolerance screening of *Helianthus annuus* seeds (Li et al., 2020). Five replicates per maternal line were used for each level of salt treatment, with a different salt treatment in each quadrant of the petri dish. Once in the growth chamber, seeds were checked every 2 days for germination. The experiment ran for 20 days. Seeds in the medium and high salt treatments failed to germinate, and were excluded from all analyses. Thus there were two treatments, control and 75 mM NaCl.

### Greenhouse experiment

Seeds were harvested in a similar manner to the germination experiment from five collection sites in Baltimore (Figure 1; Table S2). Once again, we collected *S. media* and *L. purpureum*, and also collected *V. arvensis*. We collected 12 lines from each collection environment within each collection site, which resulted in 120 lines per species and 360 lines total. All seeds were collected between April and May 2024 and set to cold stratify at 4 °C for a month.

*S. media* seeds were placed into a CARON model 7305-22-1 growth chamber, and *V. arvensis* and *L. purpureum* seeds were placed into a Percival Model I-33VL controlled environmental chamber (Percival Scientific, Perry, Iowa). Seeds were placed in plastic petri dishes with a 7.62 centimeter (3 inch) diameter on a paper towel medium, which was moistened with deionized water as needed. *S. media* seeds were grown with a 12 hr photoperiod at 20 °C/10 °C at 50% relative humidity (Roberts & Lockett, 1975), and *V. arvensis* and *L. purpureum* seeds were grown with a 12 hr photoperiod at 15 °C/6 °C with 50% relative humidity (Baskin & Baskin, 1983, 1984). *S. media* failed to germinate in sufficient quantities, so it was discarded and not included in this experiment. We began *L. purpureum* germination once *V. arvensis* had finished germinating due to limited growth chamber space. Both species grew for 30 days in the growth chamber before transfer to the Johns Hopkins University Homewood Greenhouse. This greenhouse is a heated glasshouse with an evaporative cooler and north-south light exposure, controlled with the Wadsworth STEPsaver Software Package (Wadsworth Control Systems, Arvada, Colorado). Because most of the experiment ran in late summer and early fall, we did not regulate temperature until we activated the greenhouse lights. On October 25^th^, to compensate for decreasing day lengths, we activated the greenhouse’s lighting system with a 12 hr photoperiod to provide more light. For *V. arvensis*, we planted eight maternal lines for each collection environment × site group (e.g. roadside collection environment, Patterson Park), with four replicates per maternal line per treatment (total n per experimental group = 160; Table S2). For *L. purpureum*, we planted up to eight lines for each collection environment × site group, with up to four replicates per line (Table S2). Since *L. purpureum* was less common along the roadside than the inner park, we had fewer replicates for roadside lines (n = 72) than park lines (n = 173; Table S2), but we balanced replicates between salt and control treatments for each line.

Upon transfer to the greenhouse, F1 seedlings were planted in 6 cm^3^ pots filled with 95% potting mix (The Scotts Miracle-Gro Company, Marysville, Ohio) and 5% Garden Safe brand diatomaceous earth (Spectrum Brands, Inc., St. Louis, Missouri) by volume. 24 pots, separated by treatment level and species, were randomly placed on 37 24 cm by 36 cm trays, which were rotated weekly to mitigate positional effects. Plants were given 4 weeks to acclimate to the greenhouse and were thinned to one individual per pot at the beginning of treatment. We then conducted a 2 × 2 fully factorial experimental design with two collection environments (roadside or park) and two water treatments: control (0 mM NaCL) or salt (50 mM NaCl for *L. purpureum* or 100 mM NaCl for *V. arvensis*). Salinity treatments were based off an agricultural study, where treatments of 50 or 100 mM NaCl solution revealed salinity tolerance differences between between *Capsicum annuum* lineages without causing immediate mortality (Chartzoulakis & Klapaki, 2000). All plants were bottom-watered with tap water weekly on Mondays. On Fridays, each plant in the control group was top-watered with 25 mL of tap water while each plant in the salt group was top-watered with 25 mL of saline solution. We began salt treatment for *V. arvensis* on August 20^th^, 2024, and began salt treatment for *L. purpureum* on October 11^th^, 2024. The salt treatment for *V. arvensis* was 25 mL of 50 mM NaCl in the first week to avoid environmental shock, then 25 mL of 100 mM NaCl in following weeks. We planned to use 25 mL of 100 mM NaCl for *L. purpureum*, but kept the treatment at 25 mL of 50 mM NaCl due a more adverse reaction to salt treatment arising in this species (unpublished observation). Salt treatment lasted 8 weeks for *V. arvensis* and 5 weeks for *L. purpureum*, and was stopped once salt treatment caused heavy damage or mortality in over 75% of the replicates for a species. Once lights were turned on, we added an additional weekly bottom-watering on Wednesday to compensate for greater evaporation. We concluded the experiment on November 8^th^ for *V. arvensis* and on November 15^th^ for *L. purpureum* due to widespread mortality.

For both species, we recorded plant condition weekly on a scale of 0-3, with 3 representing an undamaged plant, 2 representing a plant that sustained some damage from any source (e.g. salt burn), 1 representing a heavily damaged plant (alive with over 50% tissue damage), and 0 representing a plant that had died.

### Analyses

We analyzed data in R (R Core Team, 2024) using tidyverse (Wickham et al., 2019), and created figures using ggplot2 (Wickham et al., 2016), ggpubr (Kassambara, 2026), and cowplot (Wilke, 2025). We used an alpha of 0.05. To compare the conductivity of roadside and park soils, we used a two-way analysis of variance to determine if salinity varied by site or collection environment (roadside or park).

For our germination experiment, we calculated germination rate of the five seeds in each treatment for each maternal line. We ran linear mixed effects models using PROC MIXED in SAS (The SAS Institute, 2023) with salt treatment and collection environment as fixed effects and maternal line as a random effect. We ran separate models for each species testing for the impact of salt treatment and collection environment on germination rates. We also calculated the overall germination rate of plants in the salt treatment relative to overall germination rate of plants in the control treatment for both park and roadside plants.

In our greenhouse experiment, we assessed survival rates and plant condition. Because adverse greenhouse conditions caused high mortality in all groups at the end of the experiment, we calculated survival rates on day 51 of salt treatment for *V. arvensis* and day 29 of salt treatment for *L. purpureum*. We used PROC GLIMMIX in SAS with salt treatment and collection environment as fixed effects and maternal line as a random effect specifying a binomial distribution. To compare plant condition, we ran a repeated measures mixed model in SAS using PROC MIXED with salt treatment, collection environment, and day of experiment as fixed effects, maternal line as a random effect, and day as a repeated effect. The two species were analyzed separately. In cases of statistical significance, we used LSMeans in SAS for post-hoc analyses.

## Results

### Soil salinity

Roadside soils had significantly higher conductivity overall than park soils (F_1,48_ = 13.557, p < 0.001; Figure 1b). This pattern was consistent across collection sites, as there was no significant effect of site (F_4,48_ = 1.510, p = 0.214) or significant interaction between collection environment and site (F_4,48_ = 1.375, p = 0.257).

### Germination experiment

For *L. purpureum* we found a significant effect of salt treatment and collection environment on germination rates (Table 2) but no significant interaction. Germination rates were lower when seeds were exposed to salt treatment, and higher for park-collected lines (Figure 2). While not statistically significant, we observed a trend where the germination rate of roadside *L. purpureum* maternal lines were less negatively impacted by salt treatment (salt treatment reduced germination rate by 33%) compared with park maternal lines (salt treatment reduced germination rate by 69%) (Figure 2a, Table 2). For *S. media* we found no main effects of salt treatment or collection environment on germination rates (Figure 2a, Table 2), though *S. media* roadside line germination also appeared less impacted by salt treatment (salt treatment reduced germination rate by 60%) compared to park lines (salt treatment reduced germination rate by 83%).

**Figure 2:**
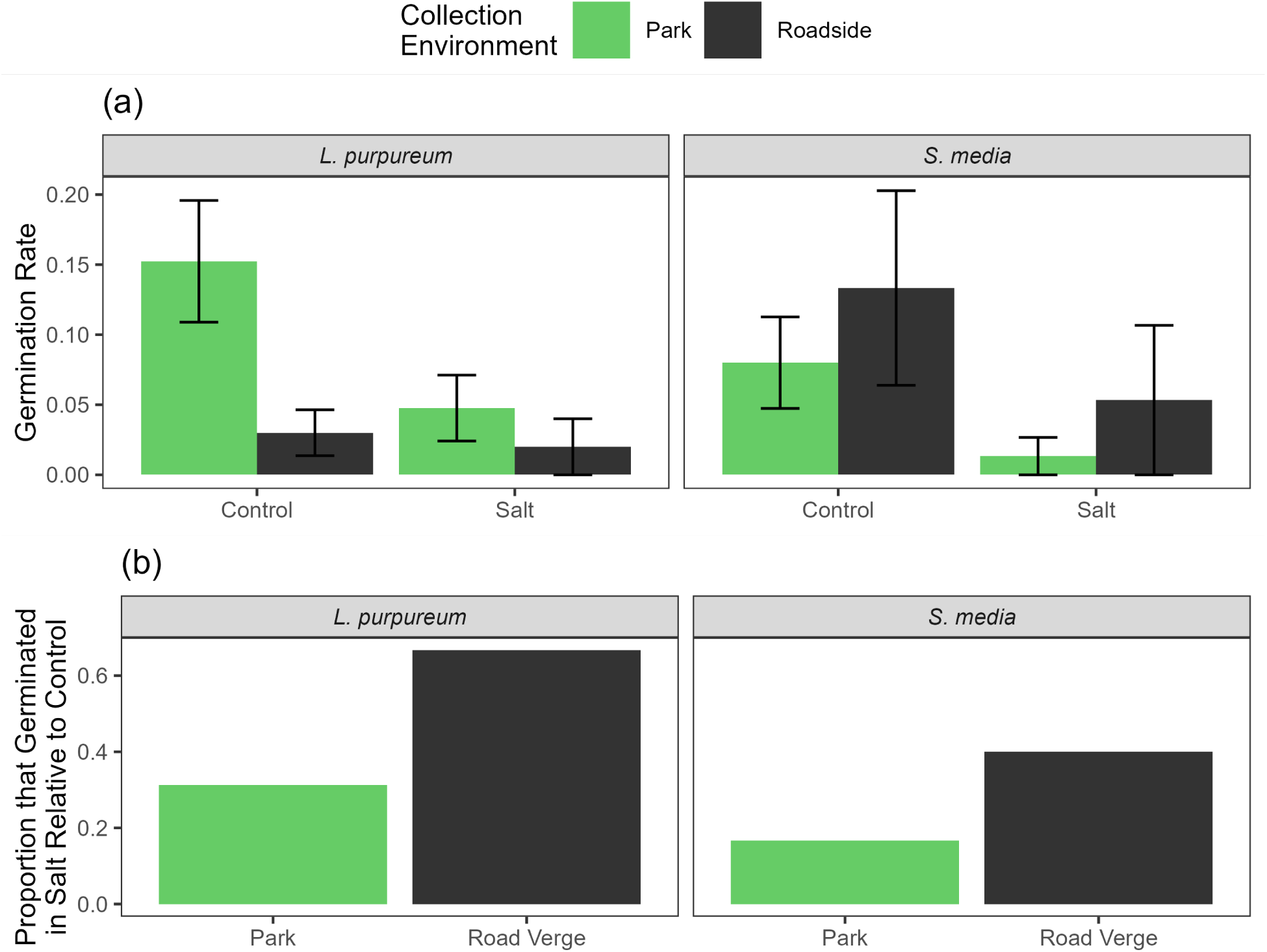
(a) Germination rates of each species; shown are means and standard error. (b) Proportion of seeds that germinated in the salt treatment group, compared with the control group.

**Table 2:**
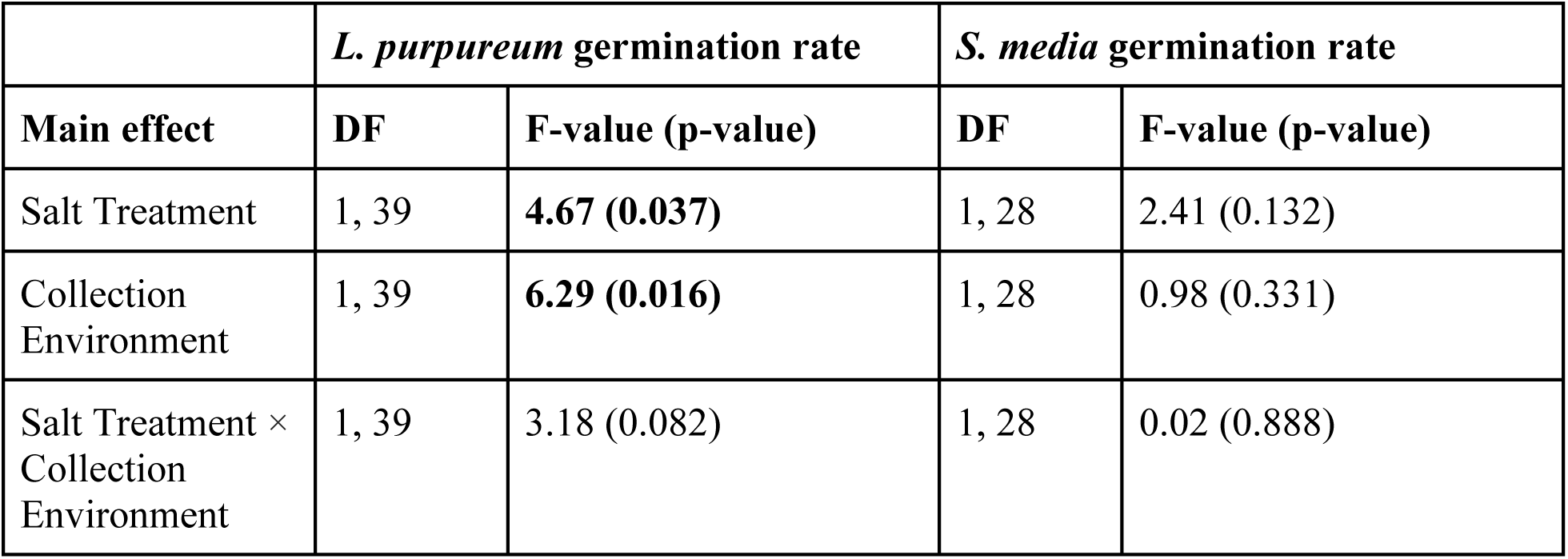
Table of main effects on germination rate of *L. purpureum* and *S. media* in the germination experiment.

|  | ***L. purpureum*** **germination rate** |  | ***S. media*** **germination rate** |  |
|---|---|---|---|---|
| **Main effect** | **DF** | **F-value (p-value)** | **DF** | **F-value (p-value)** |
| Salt Treatment | 1, 39 | **4.67 (0.037)** | 1, 28 | 2.41 (0.132) |
| Collection Environment | 1, 39 | **6.29 (0.016)** | 1, 28 | 0.98 (0.331) |
| Salt Treatment × Collection Environment | 1, 39 | 3.18 (0.082) | 1, 28 | 0.02 (0.888) |

### Greenhouse experiment

In the greenhouse experiment, we found that *V. arvensis* exposed to salt had a significantly lower survival rate (27% on day 51) than the control group (77% on day 51; Figure 3a, Table 3), but survival rate did not differ significantly by collection environment and there was no interaction. For *V. arvensis,* we found a significant three-way interaction between day of experiment, collection environment, and salt treatment on condition (Table 4). At the beginning of the experiment, plant health was similar across treatments, and the average condition of salt-treated plants worsened faster than water-treated plants (Figure 3b). After five weeks of salt treatment, park lineages had a significantly worse reaction to salt treatment than roadside lineages (Figure 3b). Towards the end of the experiment, high rates of death caused the road and park subsets of the salt group to reconverge. Around day 60 of the experiment, a fungal outbreak infected most plants across treatments, causing a steep decline in condition for all plants.

**Figure 3:**
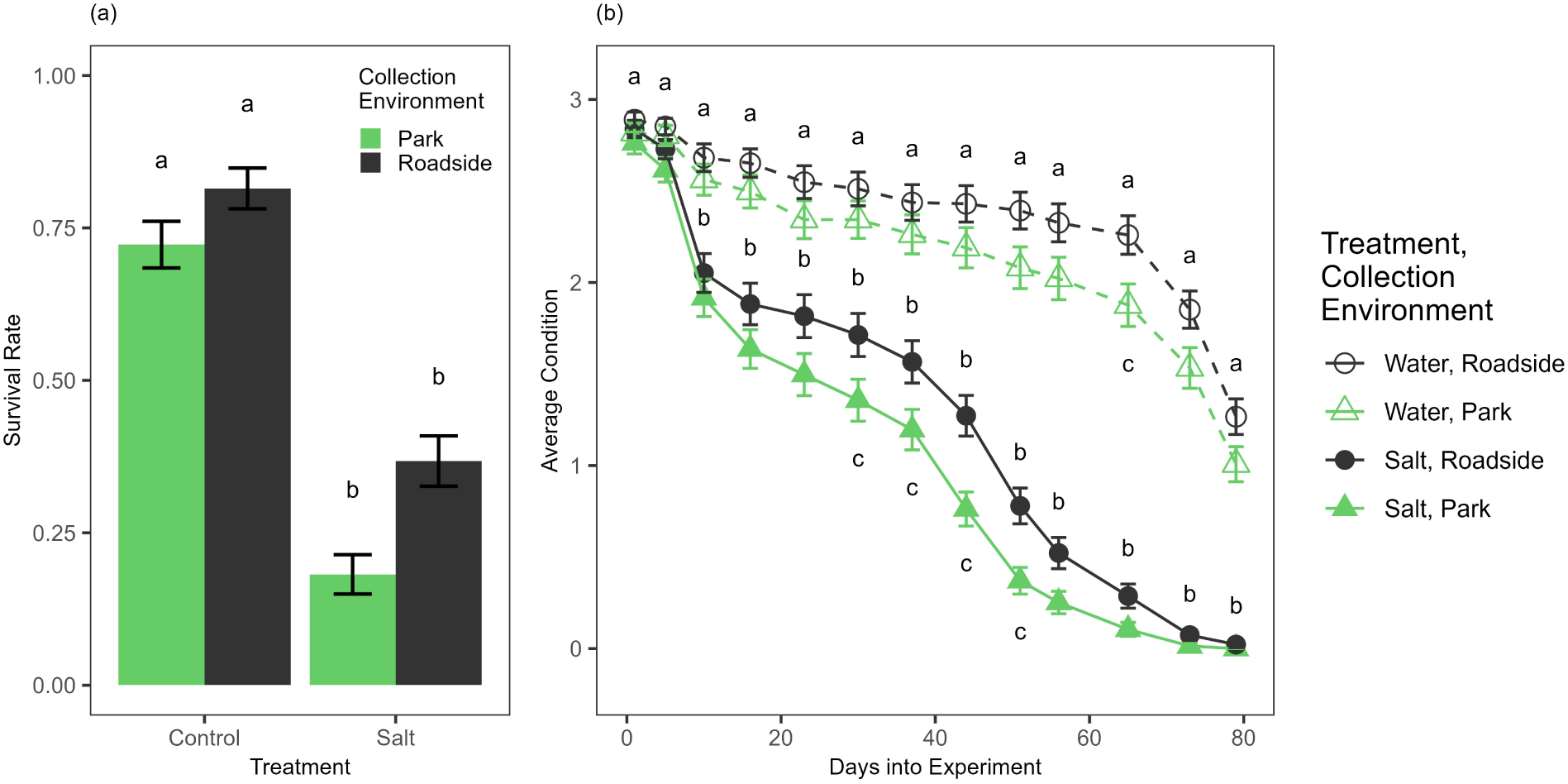
(a) *V. arvensis* survival rate on day 51 of the experiment. (b) *V. arvensis* health through the experiment by experimental group. Shown are means ± standard error; different letters indicate statistically significant differences between groups (in panel b, letters represent statistically different groups within a given date).

**Table 3:** Table of main effects on survival rate of *V. arvensis* and *L. purpureum* in the greenhouse experiment.

| **Main effect** | ***V. arvensis* survival rate** |  | ***L. purpureum* survival rate** |  |
|---|---|---|---|---|
|  | **DF** | **F-value (p-value)** | **DF** | **F-value (p-value)** |
| Salt Treatment | 1, 467 | **39.64 (<0.001)** | 1, 171 | 0.09 (0.7602) |
| Collection Environment | 1, 467 | 1.19 (0.2756) | 1, 171 | 1.60 (0.2079) |
| Salt Treatment × Collection Environment | 1, 467 | 0.20 (0.6586) | 1, 171 | 0.07 (0.7905) |

**Table 4:** Table of main effects on condition for *V. arvensis* and *L. purpureum* in the greenhouse experiment.

| **Main effect** | ***V. arvensis* condition** |  | ***L. purpureum* condition** |  |
|---|---|---|---|---|
|  | **DF (num, den)** | **F-value (p-value)** | **DF (num, den)** | **F-value (p-value)** |
| Day | 12, 7033 | **558.05 (<.001)** | 5, 1375 | **396.16 (<.001)** |
| Salt Treatment | 1, 7033 | **344.39 (<.001)** | 1, 1375 | **6.05 (0.014)** |
| Day × Salt Treatment | 12, 7033 | **101.46 (<.001)** | 5, 1375 | **3.11 (0.009)** |
| Collection Environment | 1, 7033 | 2.66 (0.103) | 1, 1375 | 1.58 (0.210) |
| Day × Collection Environment | 12, 7033 | **2.83 (0.007)** | 5, 1375 | **3.13 (0.008)** |
| Salt Treatment × Collection Environment | 1, 7033 | 0.03 (0.871) | 1, 1375 | 0.08 (0.780) |
| Day × Salt Treatment × Collection Environment | 12, 7033 | **2.09 (0.015)** | 5, 1375 | 0.68 (0.642) |

In contrast with *V. arvensis, L. purpureum* condition did not display a three-way interaction between day, treatment, and collection environment, and its survival rate did not differ by salt treatment or collection environment (Tables 3 and 4, Figure 4a). However, *L. purpureum* condition did exhibit an interaction between day and treatment (Table 4) and day and location (Table 4). Due to growth chamber space constraints, *V. arvensis* germination and growth began before *L. purpureum*, meaning that fungal infestation arose earlier in *L. purpureum’s* experimental schedule (around day 30) than for *V. arvensis*. Once again, groups diverged in the middle section of the experiment (Figure 4b, Figure 4c), where enough time had passed to identify differences but conditions in the greenhouse had not worsened enough to cause mass death.

**Figure 4:**
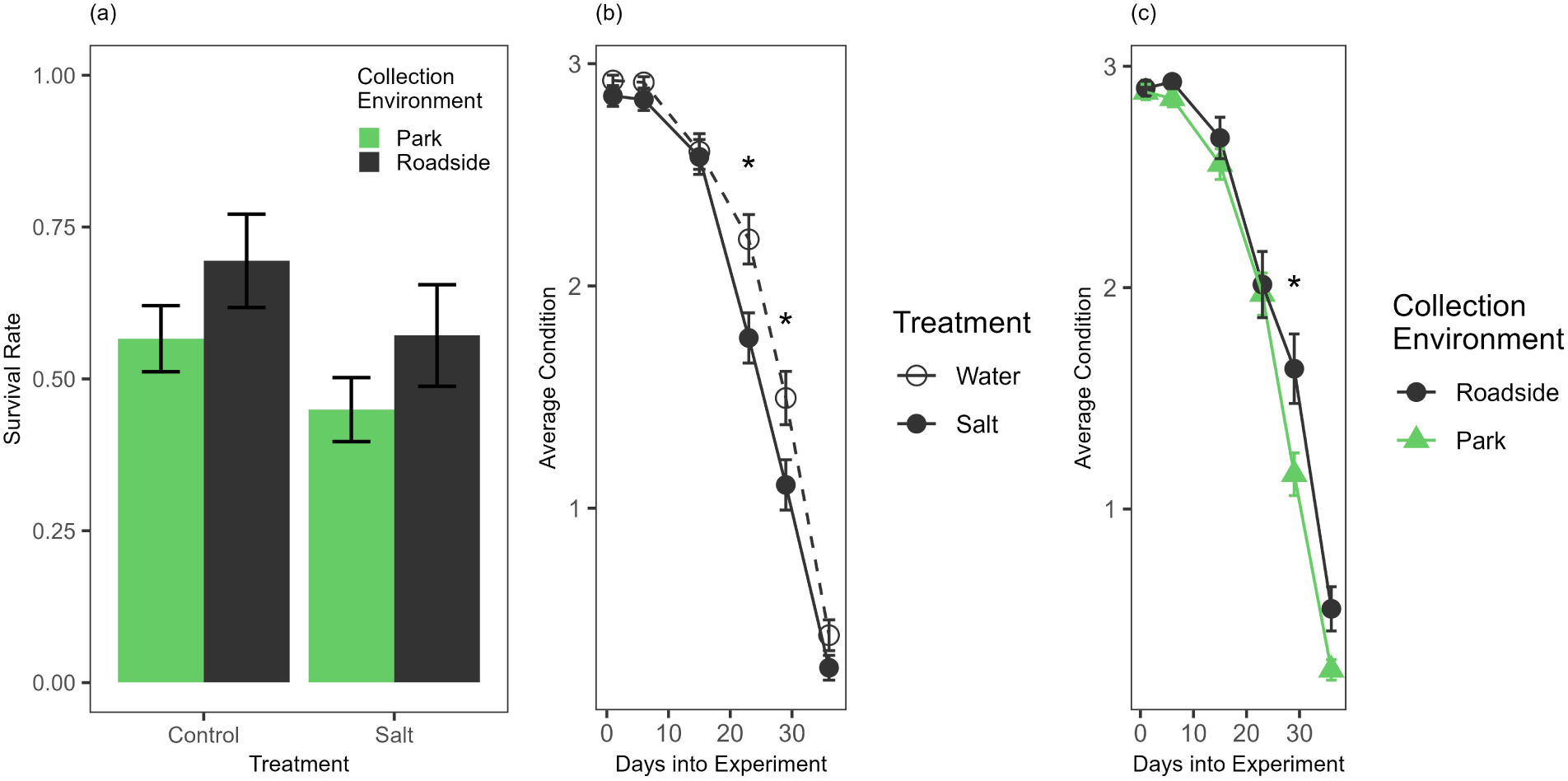
(a) *L. purpureum* survival rate on day 29 of the experiment. (b) *L. purpureum* health throughout the experiment differentiated by treatment and (c) *L. purpureum* health throughout the experiment differentiated by collection environment. Shown are means ± standard error. Asterisks indicate statistically significant differences between groups (in panels b and c, letters represent statistically different groups within a given date).

## Discussion

Roadside plants experience higher soil salinity in regions where road salt is applied in the winter, and elevated chloride pollution from deicing salts is tied to impervious surface cover (Moore et al., 2020). We studied the impact of soil salinity on the growth and survival of common urban weeds, and hypothesized that plant lineages that originated from roadsides would tolerate salt better than plant lineages that originated in co-located parks due to selection pressure and maternal effects. Importantly, we confirmed that roadside soil was indeed saltier than park soils early in the growing season. In line with common plant reactions to salinity (Łuczak et al., 2021), experimental groups exposed to salt had lower germination rates, worse condition, and lower survival rates than their clean-water counterparts. Roadside plants exhibited differences from park populations. In the germination experiment, roadside-collected *L. purpureum* seeds had significantly lower germination rates than park-collected seeds, and in the greenhouse experiment, roadside plants for both species exhibited decreased tissue damage than park plants across both treatments. Though we found some support for increased salinity tolerance in roadside-collected lines compared with park collected lines, patterns of salt tolerance varied between species and between experiments. In our germination experiment, we only observed a non-significant trend of an interaction between salt treatment and collection environment in *L. purpureum*, and did not observe an interaction between salt treatment and collection environment in *S. media*. In our greenhouse experiment, *V. arvensis* collected from the roadside displayed a significantly smaller decrease in condition compared to park-collected lines in response to salt treatment, but we observed no difference in response to salt treatment in *L. purpureum* by collection environment.

For *S. media* in the germination experiment, germination rates did not differ by experimental group. In our greenhouse experiment, only 11 of our 120 *S. media* maternal lines germinated, and two or fewer seeds germinated for each maternal line. *S. media* generally performs well in urban environments, even exhibiting increased growth in the face of moderate urban stressors (Salinitro et al., 2021). Therefore, it is likely that *S. media’s* low germination rate for the greenhouse experiment was due to unfavorable germination conditions.

We found that roadside *V. arvensis* lineages exhibited elevated salt tolerance compared to park lineages, indicative of a plastic and potentially adaptive response to saline roadside environments. This is consistent with elevated salt tolerance in roadside populations identified in *Dipsacus fullonum* subsp. *sylvestris* (Beaton & Dudley, 2004) and *Ambrosia artemisiifolia* (Eom et al., 2013), and provides further evidence that the environmental stressors of the roadside drive phenotypic differences in fast-reproducing plant populations. Because the greenhouse experiment was on the F1 generation, we are unable to tease apart whether salt tolerance differences are caused by maternal effects or genetic changes. Maternal effects can considerably impact the phenotype of plants in the following generation (Roach & Wulff, 1987), and have been specifically observed to improve the salt tolerance of plants collected from more saline habitats (El-Keblawy et al., 2016; Gu et al., 2024). Thus, we are unable to definitively conclude that adaptive evolution is occurring in *V. arvensis*. However, though they are not the gold standard of the F2 generation, F1 common garden experiments are still considered moderately strong evidence for genetic change (Lambert et al., 2021), and we take these results as tentative evidence for salt tolerance adaptation in roadside populations of *V. arvensis*.

In contrast, *L. purpureum* did not exhibit strong signs of phenotypic plasticity or adaptation to salt stress. In the germination experiment, we observed only a non-significant interaction where roadside lines had a marginally lower reduction in germination rate when exposed to salt treatment compared to park lines. In the greenhouse experiment, we observed no difference in salt tolerance between roadside and park populations. *L. purpureum* was planted a month after *V. arvensis* in the greenhouse experiment, exposing it to unfavorable conditions earlier in its salt treatment schedule than *V. arvensis* and potentially overwhelming any treatment effects. Beaton & Dudley (2010) did not observe differences in salt tolerance between roadside and park lineages of *Dipsacus fullonum* subsp. *sylvestris* in a year of severe drought, despite observing differences in this species in other years (Beaton & Dudley, 2004, 2007). However, it is also possible that urban populations of *L. purpureum* and *V. arvensis* have different reactions to elevated salt content given the variability of plant reactions to salinity (Munns & Tester, 2008).

We expected to find similar responses to salinity in our study species given that they are annual non-native species common along roadsides with similar phenology, such as early spring flowering. However, we found variation in salt tolerance among our species. Other studies of tolerance to roadside soil salinity in widespread fast-reproducing plants also identify different salt tolerance ability depending on the species. One study of *Ambrosia artemisiifolia* found that park-collected seeds exhibited reduced growth under low salinity levels, while roadside-collected seeds exhibited slightly increased growth under low salinity levels (Eom et al., 2013). Another study on the F1 generation that examined Na^+^ and Mn^2+^ tolerance in roadside weedy species found that roadside *D. fullonum* subsp. *sylvestris* lineages displayed elevated Na^+^ tolerance during germination, but did not find similar effects in *Asclepias syriaca* or *Hesperis matronalis* (Beaton & Dudley, 2004). In a follow-up study of salt tolerance in *D. fullonum* subsp. *sylvestris*, elevated seed salt concentration affected the osmotic salt tolerance of some roadside lineages, as leaching seeds before salt exposure caused these lines to lose their elevated salt tolerance, while the salt tolerance of other maternal lines did not change with leaching (Beaton & Dudley, 2007). This indicates that salt tolerance in roadside populations can be influenced by both genetic and environmental mechanisms, including developmental plasticity and maternal effects, and that these mechanisms can vary for individuals within a population. Our results contribute to a broader body of work indicating that roadside weeds exhibit elevated tolerance to heightened roadside soil salinity, that this tolerance has some heritable basis, and that tolerance to this stress varies between species. However, the relative contributions of maternal effects and genetic change to the salt tolerance of roadside lineages remains unclear, and presents an area for further study.

In our greenhouse experiment, we observed that roadside *V. arvensis* and *L. purpureum* lineages exhibited less tissue damage. The roadside is a harsher environment for plants due to many reasons beyond road salt. Lead, which is toxic to most species, was found to be over 300% more concentrated in Baltimore’s roadside compared to other urban soils (Coates et al., 2025). Elevated heat (Woudstra et al., 2024), a drier microclimate (Whitlow et al., 1992), and mechanical stress due to trampling (Oldén et al., 2021) also create adverse conditions for plants, which may favor ruderal species compared to less developed areas like meadows (Oldén et al., 2021). Accordingly, elevated tolerance of stressors besides salinity has been observed in roadside lineages of other species. For example, roadside populations of *D. fullonum* subsp. *sylvestris* were found to exhibit increased drought tolerance independently of salt tolerance (Beaton & Dudley, 2013), and elevated tolerance to soil lead content has been observed in roadside *Plantago lanceolata* lineages (Wu & Antonovics, 1976). Plants can exhibit different reactions to co-occurring stresses compared to stresses in isolation (Jiang et al., 2025), and depending on the type and intensity of the abiotic stress, combinations of abiotic stress can produce more negative or less negative effects on plants (Suzuki et al., 2014). Salt and drought stress, for example, both elicit stomatal closure to prevent excess water loss (Jiang et al., 2025), and moderate soil salinity has been observed to improve drought resistance in tomato (*Solanum lycopersicum*) (Yang et al., 2020), though this effect can vary by plant species. We speculate that the assortment of roadside stresses could cause selection pressure for generally stress-tolerant individuals.

The difference in salt tolerance and general fortitude of our roadside-collected lines compared to our park-collected lines, which were collected a short distance away from each other, emphasizes that classifying urban environments as one large habitat is limiting. It also speaks to the importance of cataloging conditions of study sites along the urban-rural gradient, as characteristics of urbanization do not often change linearly with distance from the city center (Cadenasso et al., 2007). Previous work has identified different vernalization requirements for flowering in *Taraxacum officinale* lineages collected from different urban locations, where lines collected from urban streets (over 75% impervious surface within 5 m of the plant) exhibited the highest flowering rate after low vernalization exposure compared to other urban subhabitats, potentially reflective of selection pressure for reduced vernalization (Woudstra et al., 2024). Another study identified that cyanogenesis micro-clines run in the opposite direction of broader urban-rural cyanogenesis clines (Santangelo et al., 2022b). Our results provide another example of phenotypic divergence and possible adaptation on a small spatial scale. This is especially prominent given that our focal species are capable of insect pollination (Boutin & Harper, 1991; Salehi et al., 2019). Considering that the roadside and park collection environments of the same site were rarely more than a few hundred meters away from the other, which is within the range of common North American pollinators (Hagler et al., 2011), these phenotypic responses took place at a scale where cross-pollination could still occur, further suggesting that intense heterogeneity results in different selection pressures. Future investigation into trait variation of plants and other sessile organisms between different subhabitats in close physical proximity would provide additional insight into the complex selection pressures of the urban environment.

Given the potential for roadsides to act as an invasion corridor for salt-tolerant species (Nováková et al., 2019), and that rapid adaptation is a common characteristic of successful invasive populations (Lee, 2002), urban roadsides may provide a pathway for populations of invasive species that can successfully adapt to elevated salinity levels, and future studies should evaluate the potential of saline roadsides as an avenue for invasion. Overall, we conclude that we observed tentative evidence of adaptation to higher salinity in roadside soils compared with park soils, and that salinity of the roadside environment may be a strong enough pressure to result in selection for increased salt tolerance.

## Statements & Declarations

### Funding

This project was funded by the Provost’s Undergraduate Research Award from the Johns Hopkins University Office of Undergraduate Research, and by the Hendler Global Change Award from the Morton K. Blaustein Department of Earth and Planetary Sciences.

### Competing Interests

The authors declare no competing interests.

### Author Contributions

M.L.A., A.H., J.M.C., K.C., and E.G.Y. designed the germination study. A.H. collected the seeds and J.M.C., K.C., and E.G.Y. were responsible for experimentation in the germination experiment. M.M.C. and M.L.A. designed the greenhouse study. M.M.C. was responsible for seed collection, experimentation, and measurements in the greenhouse experiment, in addition to soil collection and salinity measurement. M.M.C., M.L.A., and J.M.C. contributed to data analysis. M.M.C. and M.L.A. wrote the manuscript with input from all co-authors.

### Data and Code Availability

R and SAS code is available at https://github.com/maxmcarroll/Salt_Tolerance_Urban_Weeds. Data will be made publicly available at Environmental Data Initiative upon acceptance.

## Acknowledgments

We thank Dr. Katalin Szlavecz for her advice on soil collection and analysis. We also thank Vicki Li for her help with the germination experiment, and Claudia Ouimet, Mehr Khosla, and Cassidy Woodham for help in the greenhouse. Finally, we thank Ethan Gale and Diego Trafton for help procuring materials for the greenhouse experiment.

## Notes

### Competing Interest Statement

The authors have declared no competing interest.

