## Supplementary Information for "Examining urban weed response to elevated salt levels"

**Supplementary Information for the manuscript "Examining urban weed response to elevated salt levels"**

**Table S1:** Number of maternal lines grouped by species, site, and collection environment (roadside or park). For each maternal line, 5 seeds were germinated in each treatment (control, low salt, medium salt, and high salt treatments).

| **Site** | ***Stellaria media*** | | ***Lamium purpureum*** | |
|---|---|---|---|---|
| | Park | Roadside | Park | Roadside |
| Carroll Park | 3 | 3 | 1 | 0 |
| Druid Hill Park | 3 | 3 | 4 | 4 |
| Montebello/Herring Run Park | 3 | 3 | 6 | 4 |
| Patterson Park | 3 | 3 | 2 | 3 |
| Wyman Park | 3 | 3 | 8 | 9 |

**Table S2:** Number of maternal lines grouped by species, site, and collection environment (roadside or park), with number of replicates per grouping in parentheses. Each line had up to four replicates per treatment. For each of these groups, replicates were divided evenly between salt and water treatment.

| **Site** | ***Veronica arvensis*** | | ***Lamium purpureum*** | |
|---|---|---|---|---|
| | Park | Roadside | Park | Roadside |
| Clifton Park | 8 (64) | 8 (64) | 8 (41) | 7 (23) |
| Druid Hill Park | 8 (64) | 8 (64) | 6 (28) | 8 (20) |
| JHU (The Glade) | 8 (64) | 8 (64) | 8 (44) | 4 (7) |
| Patterson Park | 8 (64) | 8 (64) | 5 (14) | 4 (9) |
| Riverside Park | 8 (64) | 8 (64) | 8 (46) | 3 (13) |

**Table S3:** Average electrical conductivity (EC) for each site, grouped by site and collection environment.

| Site | Park EC (microSiemens) ± SE | Park Soil Sample Count | Roadside EC (microSiemens) ± SE | Roadside Soil Sample Count |
|---|---|---|---|---|
| Clifton Park | 53.783 ± 6.006 | 6 | 155.960 ± 68.550 | 5 |
| Druid Hill Park | 44.083 ± 5.913 | 6 | 163.667 ± 17.985 | 6 |
| JHU (The Glade) | 63.360 ± 8.659 | 5 | 91.883 ± 10.917 | 6 |
| Patterson Park | 32.133 ± 3.981 | 6 | 245.93 ± 109.556 | 6 |
| Riverside Park | 65.150 ± 6.022 | 6 | 388.417 ± 166.831 | 6 |
